# Rapid Emergence of Resistance to Broad-Spectrum Direct Antimicrobial Activity of Avibactam

**DOI:** 10.1101/2024.09.25.615047

**Authors:** Michelle Nägeli, Shade Rodriguez, Abigail L. Manson, Ashlee M. Earl, Thea Brennan-Krohn

## Abstract

Avibactam (AVI) is a diazabicyclooctane (DBO) β-lactamase inhibitor used clinically in combination with ceftazidime. At concentrations higher than those typically achieved *in vivo*, it also has broad-spectrum direct antibacterial activity against *Enterobacterales* strains, including metallo-β-lactamase-producing isolates, mediated by inhibition of penicillin-binding protein 2 (PBP2). This activity is mechanistically similar to that of more potent novel DBOs (zidebactam, nacubactam) in late clinical development. We found that resistance to AVI emerged readily, with a mutation frequency of 2×10^−6^ to 8×10^−5^. Whole genome sequencing of resistant isolates revealed a heterogeneous mutational target that permitted bacterial survival and replication despite PBP2 inhibition, in line with prior studies of PBP2-targeting drugs. While such mutations are believed to act by upregulating the bacterial stringent response, we found a similarly high mutation frequency in bacteria deficient in components of the stringent response, although we observed a different set of mutations in these strains. Although avibactam-resistant strains had increased lag time, suggesting a fitness cost that might render them less problematic in clinical infections, there was no statistically significant difference in growth rates between susceptible and resistant strains. The finding of rapid emergence of resistance to avibactam as the result of a large mutational target has important implications for novel DBOs with potent direct antibacterial activity, which are being developed with the goal of expanding cell wall-active treatment options for multidrug-resistant gram-negative infections but may be vulnerable to treatment-emergent resistance.

## INTRODUCTION

Beta-lactamase inhibitors have a long history in the arms race between humans and microbes. By defending β-lactam antibiotics against bacterial β-lactamase enzymes, they extend the activity spectrum of β-lactams, which are valued as first-line therapeutic agents because of their safety, efficacy, and long clinical track record (1). Beta-lactamase inhibitors have grown increasingly important in the era of multidrug resistance, as broad-spectrum β-lactamases, including carbapenemases, have emerged as a key β-lactam resistance mechanism among gram-negative bacteria (2). For decades, all β-lactamase inhibitors in clinical use were themselves β-lactam compounds that serve as “suicide inhibitors” of β-lactamase enzymes (3). Despite their β-lactam structure, these compounds have minimal intrinsic antibacterial activity (with a few exceptions, most notably sulbactam, which is active against *Acinetobacter baumannii* (3)). In 2015, the first non-β-lactam β-lactamase inhibitor, avibactam, was approved by the FDA in a combination product with ceftazidime. Avibactam was the first β-lactamase inhibitor with activity against serine carbapenemase enzymes, including *Klebsiella pneumoniae* carbapenemases (KPCs), thus rendering ceftazidime-avibactam the first β-lactam-based agent that could be used to treat infections caused by carbapenem-resistant *Enterobacterales* (CRE).

Avibactam (AVI) is a diazabicyclooctane (DBO) compound (Figure 1). It exerts its β-lactamase inhibitor activity through covalent, reversible binding to serine β-lactamases such as *Klebsiella pneumoniae* carbapenemases (KPCs) (4), although it does not inhibit metallo-β-lactamase carbapenemases (MBLs) (5). We observed that AVI also has direct *in vitro* antimicrobial activity against *Enterobacterales* isolates, including MBL-producing strains, a phenomenon mediated by its binding of penicillin-binding protein 2 (PBP2) (6), one of the classes of transpeptidase enzymes involved in bacterial peptidoglycan synthesis that constitute the targets of β-lactam drugs (7). Despite prior reports in the literature of direct AVI activity, the drug is typically described as lacking intrinsic antibacterial activity (8, 9), probably because serum levels achieved with standard doses of ceftazidime-avibactam are unlikely to be high enough to exert significant direct activity (10) given the range of AVI MICs (8-32 μg/mL). In recent years, however, DBOs with much more potent direct antimicrobial activity have been developed. One such compound, zidebactam, is currently undergoing phase 3 trials in combination with cefepime (11, 12). The spectrum of zidebactam’s direct activity includes *Pseudomonas aeruginosa*, and case reports have described compassionate use of cefepime-zidebactam for treatment of MBL-producing *P. aeruginosa* infections (13, 14). Because zidebactam, like avibactam, does not inhibit MBL enzymes, the activity of cefepime-zidebactam against MBL-producing strains results from the intrinsic PBP2-inhibiting activity of zidebactam. (A β-lactam “enhancer” effect, in which residual combination activity may be present in DBO-β-lactam combinations resistant to each component of the combination individually, has also been described (11, 15).) As zidebactam and other more potent DBOs were not widely available at the time we undertook these investigations, we used AVI as a model compound to explore direct DBO activity. We observed a high rate of resistance mutation frequency to AVI, a phenomenon that is familiar from experience with the β-lactam mecillinam (amdinocillin), the only clinically available drug that exclusively targets PBP2 (16), and has also been observed in early investigations of nacubactam, another DBO with potent direct antibacterial activity (17).

**Figure 1.**
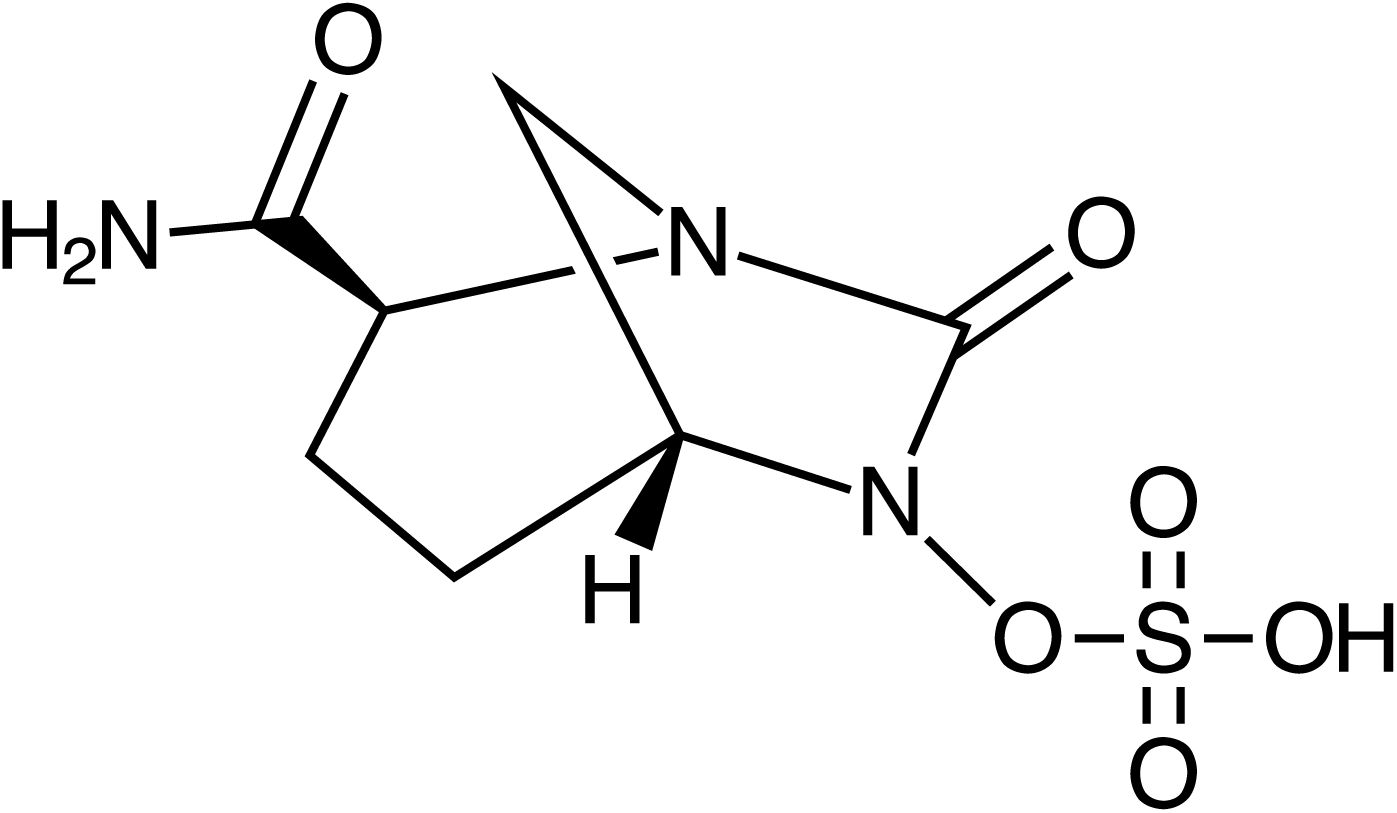
Chemical structure of avibactam

In this paper, we describe the rapid emergence of stable, high-level resistance to direct AVI activity, which appears to result from a high frequency of resistance-conferring mutations and causes cross-resistance to other PBP2-targeting drugs. We also characterize these mutations through whole-genome sequencing. As is the case with mecillinam and nacubactam, AVI resistance involves a large and heterogeneous mutational target, with mutations in different AVI-resistant strains occurring in numerous different genes. The mechanism of resistance to mecillinam and nacubactam is believed to involve upregulation of the stringent response, resulting in compensatory changes that allow bacterial cells to tolerate PBP2 inhibition (16, 18), and our sequencing results, as well as morphological analysis of bacterial cells, demonstrates the same phenomenon with AVI. Some of our findings, including fitness impairment in AVI-resistant strains as well as increased activity of AVI in an immunocompetent relative to a neutropenic mouse model, suggest that treatment-emergent resistance may be less problematic clinically than it appears *in vitro*. Overall, however, our results indicate that further study of resistance to novel DBO agents will be essential in ensuring that these drugs can be used effectively to treat patients with multidrug-resistant gram negative infections, and in particular infections caused by MBL-producing organisms, in which resistance to PBP2 inhibition may render DBO-containing β-lactam/β-lactamase inhibitor combinations ineffective.

## RESULTS

### Avibactam has broad-spectrum activity against *Enterobacterales*

Avibactam MICs of 74 gram-negative bacterial isolates, enriched for carbapenemase-producing organisms, were tested using the digital dispensing method (DDM) (Table 1, Table S1). MICs ranged from 4 to >64 μg/mL. MIC_50_ and MIC_90_ for *Enterobacterales* (n = 54) were 16 μg/mL and 64 μg/mL, respectively, while all *Pseudomonas aeruginosa* and *Acinetobacter baumannii* isolates (n = 20) had MICs of >64 μg/mL. Among isolates with low AVI MICs (8 μg/mL) were two extremely multidrug-resistant strains: *K. pneumoniae* FDA-CDC 0636 (the pan-resistant “Nevada” strain, which encodes an NDM-1 metallo-β-lactamase (19, 20)) and *E. coli* ARLG 2829 (the first strain identified in the US containing both a carbapenemase (NDM-5) and a mobile colistin resistance gene (*mcr-1*) (21)). To further investigate AVI activity against these isolates, time-kill studies were performed with AVI concentrations of 8, 16, 32, 64, and 128 μg/mL (Figure 2). At the MIC for both strains (8 μg/mL), growth was inhibited through 6 hours, but there was no significant decrease in colony count. At higher concentrations of AVI, cell counts fell by 0.5-3.3 log_10_ CFU/mL from starting inoculum by 6 hours, but then began to increase. By 24 hours, cell density had increased by 2.0-3.0 log_10_ CFU/mL from starting inoculum even in cultures treated with AVI 128 μg/mL (16x MIC), although final colony counts were lower than in the untreated growth control.

**Figure 2.**
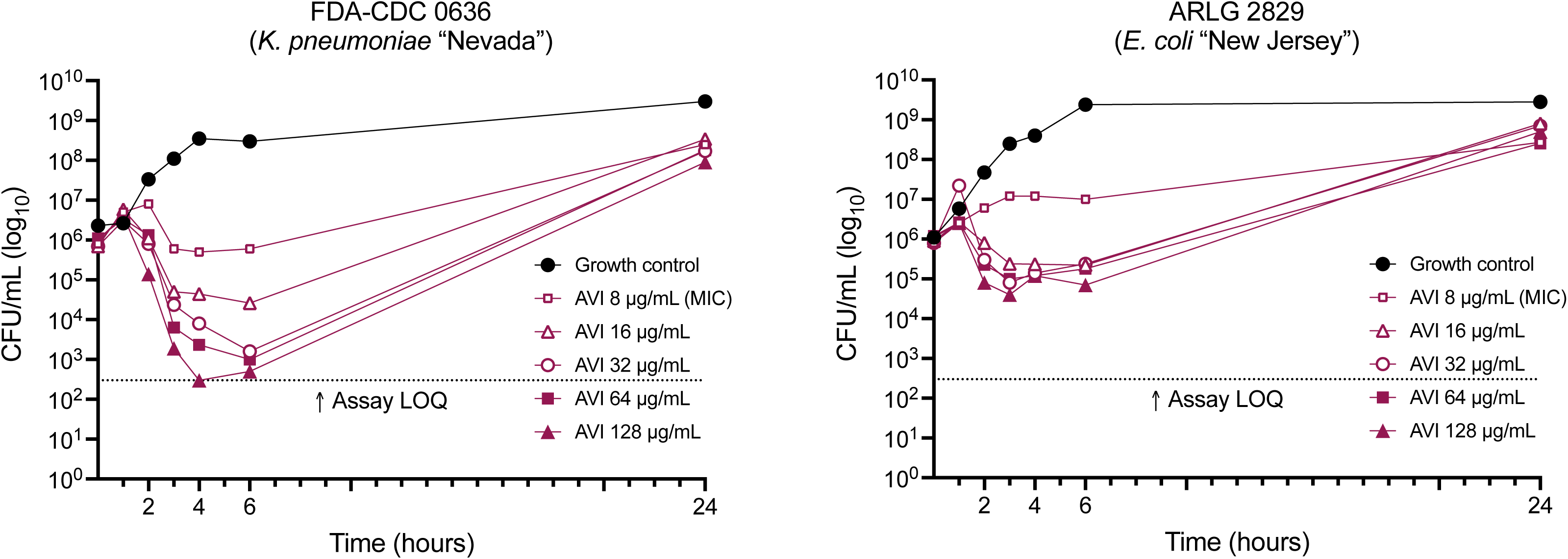
Time-kill studies of *K. pneumoniae* FDA-CDC 0636 and E. coli ARLG 2829 grown with avibactam AVI, avibactam; LOQ, limit of quantitation

**TABLE 1.** Avibactam minimal inhibitory concentrati.

| Category | MIC<sub>50</sub> | MIC<sub>90</sub><br>µg/mL | MIC range |
| --- | --- | --- | --- |
| All strains (n = 74) | 32 | 64 | 4 - >64 |
| *Enterobacterales* (n = 54) | 16 | 64 | 4 - >64 |
| *E. coli* (n = 15) | 8 | 64 | 4 - >64 |
| *Klebsiella pneumoniae* (n = 25) | 16 | 64 | 4 - >64 |
| Other *Enterobacterales* (n = 14) | 16 | 64 | 16 - >64 |
| Carbapenemase-producers (non-MBL) (n = 16) | 8 | 64 | 4 - >64 |
| MBL-producers (n = 33) | 16 | 64 | 8 - >64 |
| *Pseudomonas aeruginosa* (n = 5) | >64 | >64 | >64 |
| *Acinetobacter baumannii* (n = 15) | >64 | >64 | >64 |

MBL: Metallo-β-lactamase

### Avibactam activity is mediated by inhibition of penicillin-binding protein 2 (PBP2)

Although AVI is not a β-lactam compound, it is known to bind to and inhibit penicillin-binding protein 2 (PBP2) (6, 22). Inhibition of different PBPs induces distinct morphological changes, with PBP2 inhibition in *Enterobacterales* resulting in generation of enlarged, rounded cells (23). Serial Gram stain images of bacteria treated with AVI as in time-kill experiments were obtained under oil immersion magnification. At concentrations at and above the MIC, cells developed the distinctly rounded and enlarged appearance classically observed in bacteria treated with PBP2 inhibiting drugs (Figure 3a). Notably, bacterial cells that are resistant to PBP2 targeting drugs such as mecillinam and nacubactam exhibit rounding during treatment with the drug, even though they are still able to survive and replicate (15, 24, 25), as resistance typically involves compensatory mutations in genes other than PBP2. To assess whether the same effect occurred with AVI, bacteria were grown with 128 μg/mL AVI as in time-kill experiments, subcultured overnight on antibiotic-free media, then grown again for 24 hours in liquid culture containing AVI at 8 μg/mL and at 128 μg/mL. Although the AVI MIC, performed in parallel with the growth curve experiment, was confirmed as >256 μg/mL and the resistant cells grew to within 0.5 log_10_ CFU/mL of the untreated resistant strain by 24 hours (Figure 4), Gram stain images at 3 and 24 hours showed rounded morphology of the cells at both AVI concentrations (Figure 3b), indicating preservation of active PBP2 inhibition even in bacteria able to survive and replicate during AVI exposure.

**Figure 3.**
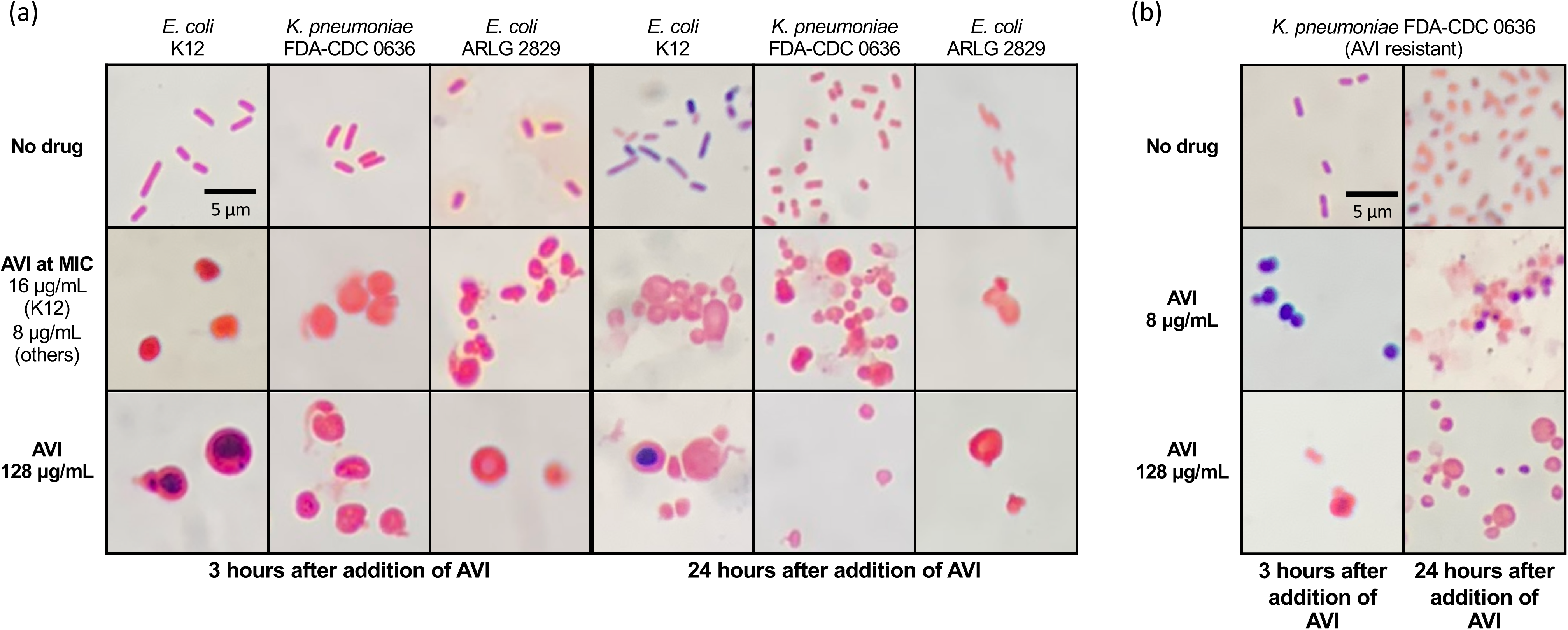
Gram stain images of avibactam-susceptible cells (a) and avibactam-resistant cells (b) grown with different concentrations of avibactam, 1000X AVI, avibactam.

**Figure 4.**
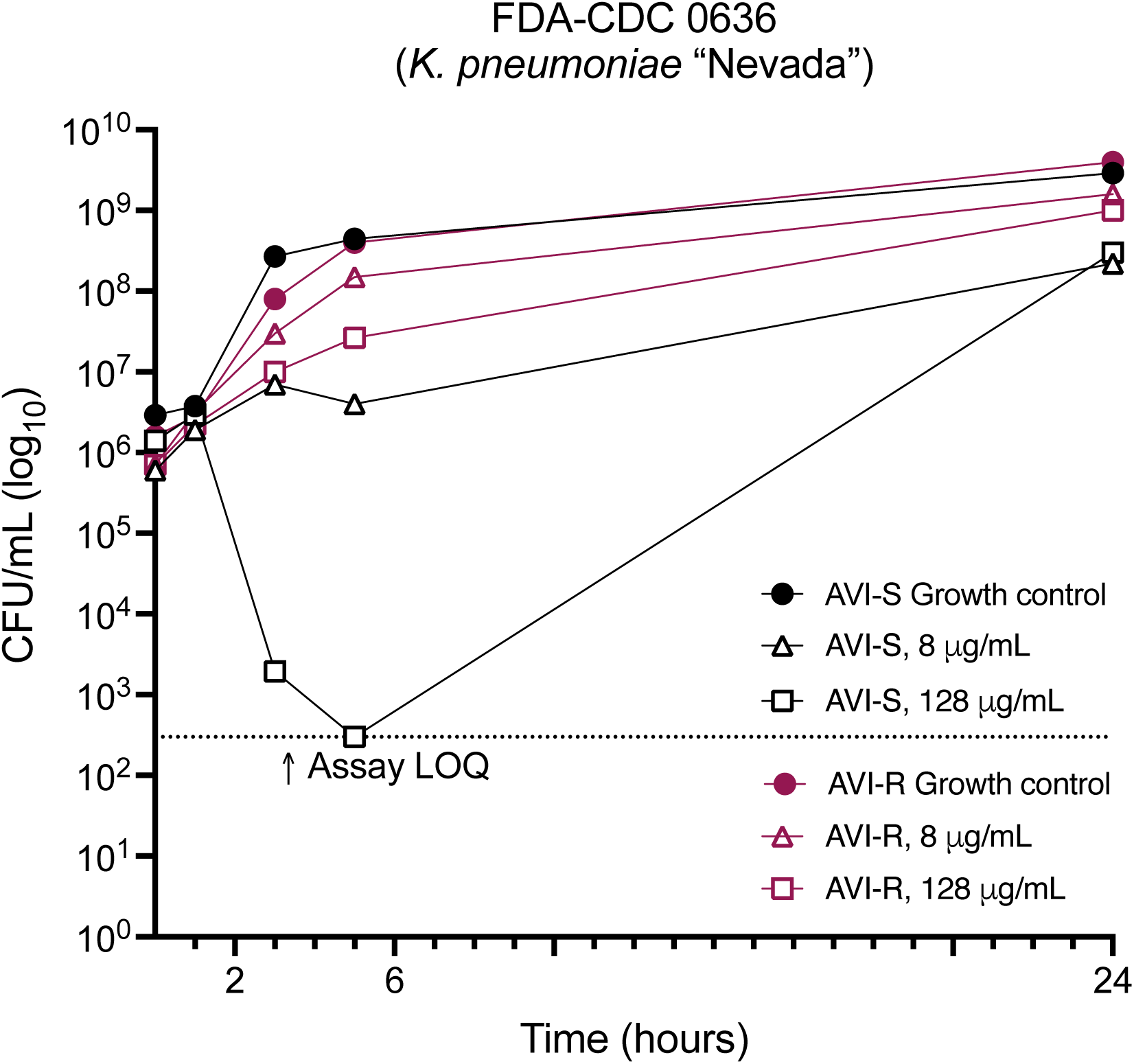
Time-kill studies of *K. pneumoniae* FDA-CDC 0636 parent and avibactam-resistant derivative strains grown with avibactam. AVI, avibactam; LOQ, limit of quantitation

Cross-resistance between AVI and other PBP-targeting drugs was assessed using two broadly β-lactam-susceptible isolates (*K. pneumoniae* BIDMC 22 and *E. coli* BIDMC 49A). Following growth for 24 hours in AVI 128 μg/mL, the MICs of mecillinam, a β-lactam antibiotic which, like AVI, exclusively targets PBP2 (22, 26), increased 16x for BIDMC 22 and >64x for BIDMC 49A, while the MIC of zidebactam, a DBO with potent PBP2-inhibiting activity (27), increased by >1024x for both strains (Table 2). In two other cases (amoxicillin for BIDMC 22 and cefepime for BIDMC 49A), MICs increased 4x; both of these drugs do exert partial PBP2 binding (28, 29). Post-AVI MICs in all others changed by no more than a single 2-fold doubling dilution, which is within the expected range of variability for MIC assays.

**TABLE 2.** Effect of AVI resistance on MICs of other drugs.

| Drug | PBP target(s) | *K. pneumoniae* BIDMC 22 |  |  | *E. coli* BIDMC 49A |  |  |
| --- | --- | --- | --- | --- | --- | --- | --- |
|  |  | Initial MIC | Post-AVI MIC | MIC change | Initial MIC | Post-AVI MIC | MIC change |
| Avibactam | 2 | 16 | 128 | **8x** | 8 | 128 | **16x** |
| Mecillinam | 2 | 0.25 | >16 | **>64x** | 0.063 | 1 | **16x** |
| Zidebactam | 2 | 0.125 | >128 | **>1024x** | 0.125 | >128 | **>1024x** |
| Meropenem | 2>4>3>1 | 0.063 | 0.063 | None | 0.016 | 0.031 | 2x |
| Amoxicillin | 4>2>3 | 8 | 32 | **4x** | 4 | 2 | -2x |
| Cefepime | 3>2>1>4 | 0.063 | 0.063 | None | 0.031 | 0.125 | **4x** |
| Ceftazidime | 3>1 | 0.25 | 0.25 | None | 0.25 | 0.25 | None |
| Aztreonam | 3 | 0.063 | 0.063 | None | 0.063 | 0.125 | 2x |

AVI: Avibactam; PBP: penicillin-binding protein. MICs are expressed in µg/mL.

### Avibactam resistance emerges readily during drug exposure and persists in the absence of selective pressure

To determine whether regrowth of bacteria in time-kill experiments was the result of the development of heritable AVI resistance, cells were recovered after 24 hours of growth in media containing 128 μg/mL AVI, and sequential AVI MIC testing was performed on isolates subjected to serial daily subcultures on antibiotic-free media over the course of 15 days. The MICs of the 3 different strains on which this procedure was performed (*E. coli* K12, *K. pneumoniae* FDA-CDC 0636, and *E. coli* ARLG 2829) remained >256 μg/mL over the course of the experiment (32-64x starting MIC). Furthermore, to confirm that drug breakdown over the course of the experiment had not contributed to regrowth during the initial time-kill experiment, a biological assay was performed to determine the approximate concentration of active AVI remaining in the supernatant at the end of a 24-hour time-kill study. The results demonstrated that the concentration remained ~128 μg/mL at the completion of the experiment.

A diffusion-based tolerance test (30, 31) was performed to determine whether tolerance to AVI was also contributing to regrowth (32). A lawn of *E. coli* K12 was incubated overnight in the presence of an AVI-impregnated disk, which was then removed and replaced with a glucose-containing disk for nutrient repletion, to allow for regrowth of colonies from any tolerant cells that had survived in the ~20 mm zone of clearance. After a subsequent night of incubation, no colonies had appeared in the zone of clearance, indicating an absence of AVI tolerance and confirming that regrowth in time-kill studies represented true heritable resistance.

### Rapid emergence of avibactam resistance is the result of a high resistance mutation frequency, even in stringent response-deficient strains

Because resistance to PBP2-targeting drugs is thought to involve activation of the stringent response (18), time-kill studies were also performed using derivatives of *E. coli* K12 with inactivating mutations in the stringent response pathway (33) as well as the SOS response pathway (34) (Table S2). Strains were grown for 48 hours with AVI at 128 μg/mL (8-16 x MIC). At 24 hours, the Δ*spoT* strain had regrown to a similar density as the K12 parent strain, while the Δ*recA* and Δ*relA* strains had cell densities that were lower by 0.8 and 1.5 log_10_ CFU/mL, respectively, although both had still regrown above the starting inoculum. The most notable effect on regrowth at 24 hours was seen in the strain lacking both *relA* and *spoT*, which had 3.4 log_10_ CFU/mL fewer cells than the parent strain and 1.8 log_10_ CFU/mL fewer than its own starting inoculum. By 48 hours, all treated strains had a similar cell density (Figure 5).

**Figure 5.**
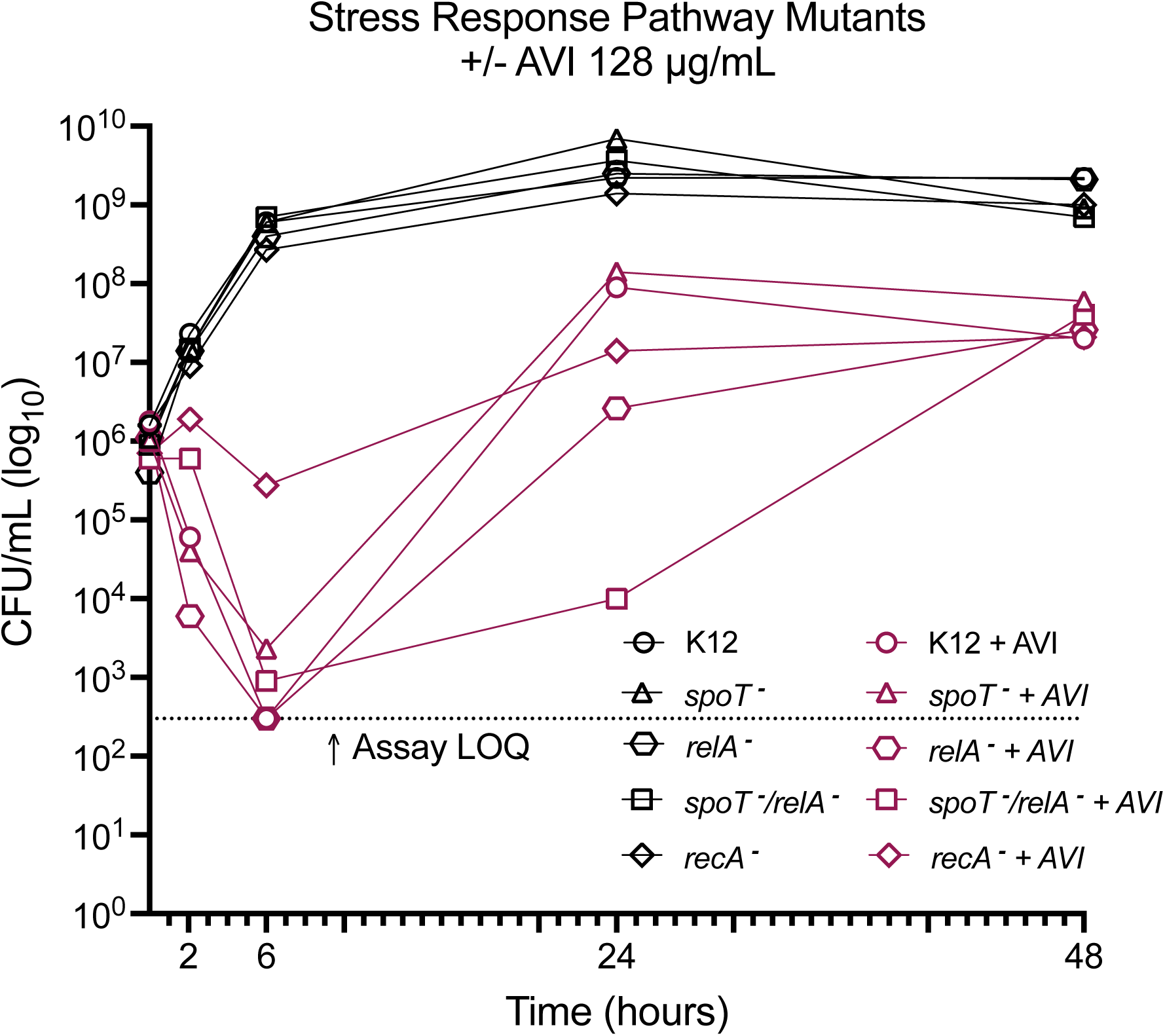
Time-kill studies of *E. coli* K12 and strains with mutations in stringent response (*spoT*, *relA*) and SOS (*recA*) genes. AVI, avibactam; LOQ, limit of quantitation

The mutation frequency for *E. coli* K12 was 8.5 ×10^−6^ and 6.6 ×10^−6^ at 4x and 8x MIC, respectively (Figure 6). Mutation frequency rates for the other strains tested were similar. Interestingly, there was no statistically significant decrease in mutation frequency between K12 and strains with deletions in key stringent response genes (p > 0.5 by unpaired two-tailed t-test); indeed, the only significant differences in mutation frequency between K12 and other strains were an increase in mutation frequency at 4x MIC in a Δ*spoT* strain (p = 0.048) and a Δ*recA* strain (p = 0.043).

**Figure 6.**
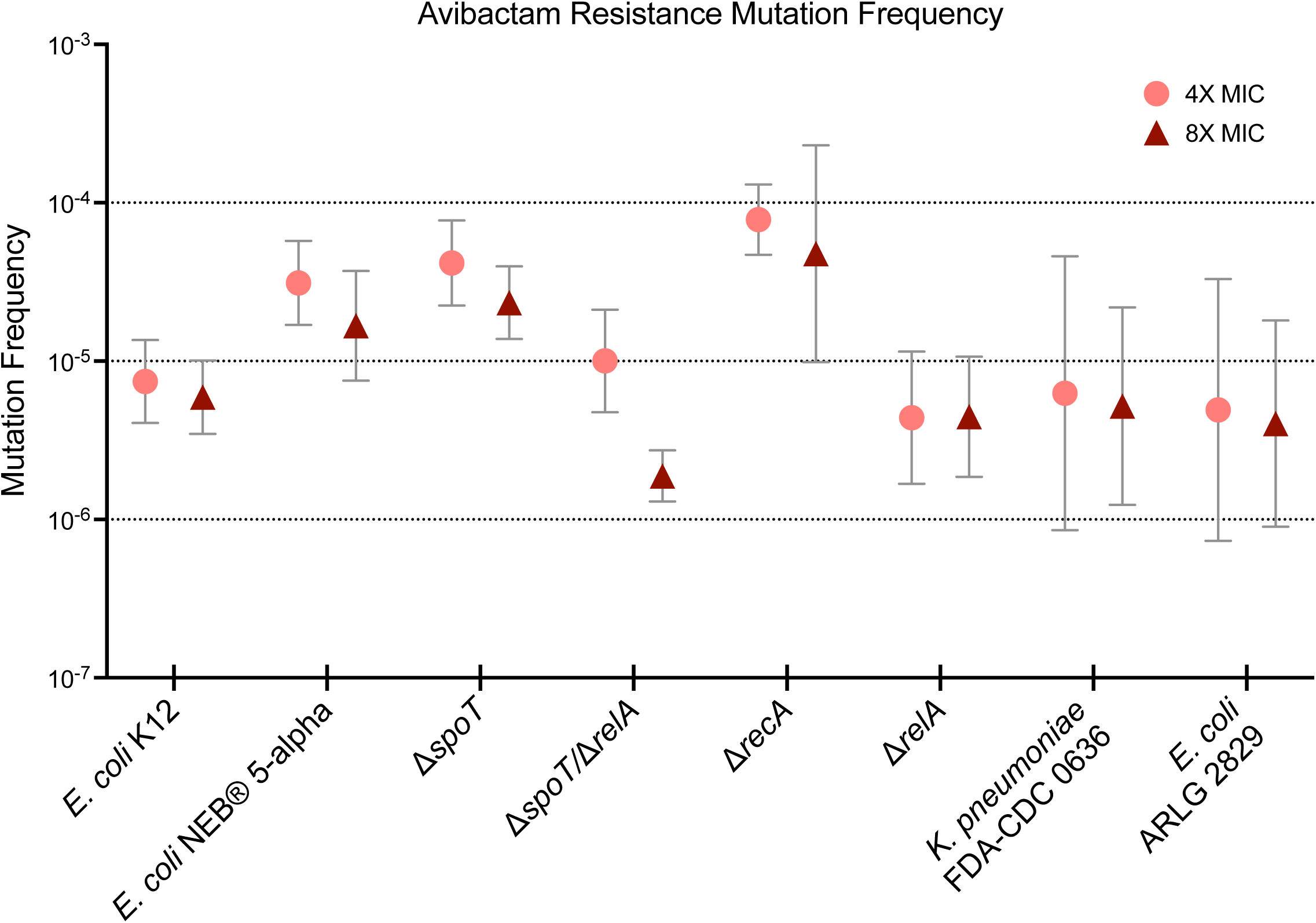
Avibactam resistance mutation frequency at multiples of the AVI MIC. Symbols and bars indicate geometric mean and standard deviation of 3 replicates.

### A diverse set of mutations and methylation changes underlie avibactam resistance

Illumina sequencing was performed on 2-3 AVI-R mutants each of *E. coli* K12, a K12 Δ*spoT* derivative, a K12 Δ*spoT*/Δ*relA* derivative, and NEB® 5-alpha in order (a) to determine whether AVI resistance-related mutations were similar to those seen in bacteria resistant to other PBP2-targeting drugs like mecillinam and nacubactam (16, 18) and (b) to determine whether absence of stringent and SOS response genes would result in a different mutational pattern. Sequencing reads were aligned to the *E. coli* K12 reference genome, and variants were called using Pilon (35) (Table 3). One of the K12 mutants had a 1336 bp insertion sequence (IS2) 145 bp upstream of threonine-tRNA ligase (*thrS*), likely in the promoter region of the gene; mutations in *thrS* have previously been reported in mecillinam-resistance *E. coli* strains (16). Both K12 mutants had point mutations causing amino acid changes in *cyaA* (adenylate cyclase), another gene that has previously been implicated in mecillinam resistance (36). None of the mutations seen in the two Δ*spoT* mutants have been previously described, but both strains had the same amino acid change (A63D) in *fabR*, which encodes a transcriptional regulator that represses unsaturated fatty acid synthesis (37). In five of the ten strains (one K12 mutant, both Δ*spoT* mutants, and 2 of 3 Δ*spoT*/Δ*relA* mutants), there was a large intergenic insertion between an IS5 family transposase and *oppA*, which encodes an oligopeptide ABC transporter periplasmic binding protein. OppA is the periplasmic component of an oligopeptide transport system and has also been implicated as a cause of aminoglycoside resistance, possibly because the protein plays a role in aminoglycoside uptake by the cell (38). One of the Δ*spoT*/Δ*relA* mutants had a 7 bp deletion resulting in a premature stop codon in *tolB*, the gene encoding a periplasmic protein in the Tol-Pal system, which is involved in bacterial cell division (39). A premature stop codon in *tolB* has previously been described in a nacubactam-resistant isolate (18). Interestingly, each of the three mutants of NEB® 5-alpha had a 1338 bp insertion within *cysB*, the gene encoding the transcriptional regulator CysB, which controls cysteine biosynthesis. Inactivating mutations in *cysB* have been found in the majority of clinical mecillinam-resistant isolates, potentially because they cause a lower fitness cost than other mutations conferring resistance to PBP2 inhibition, yet have rarely been reported in laboratory-selected strains (16).

**TABLE 3.**
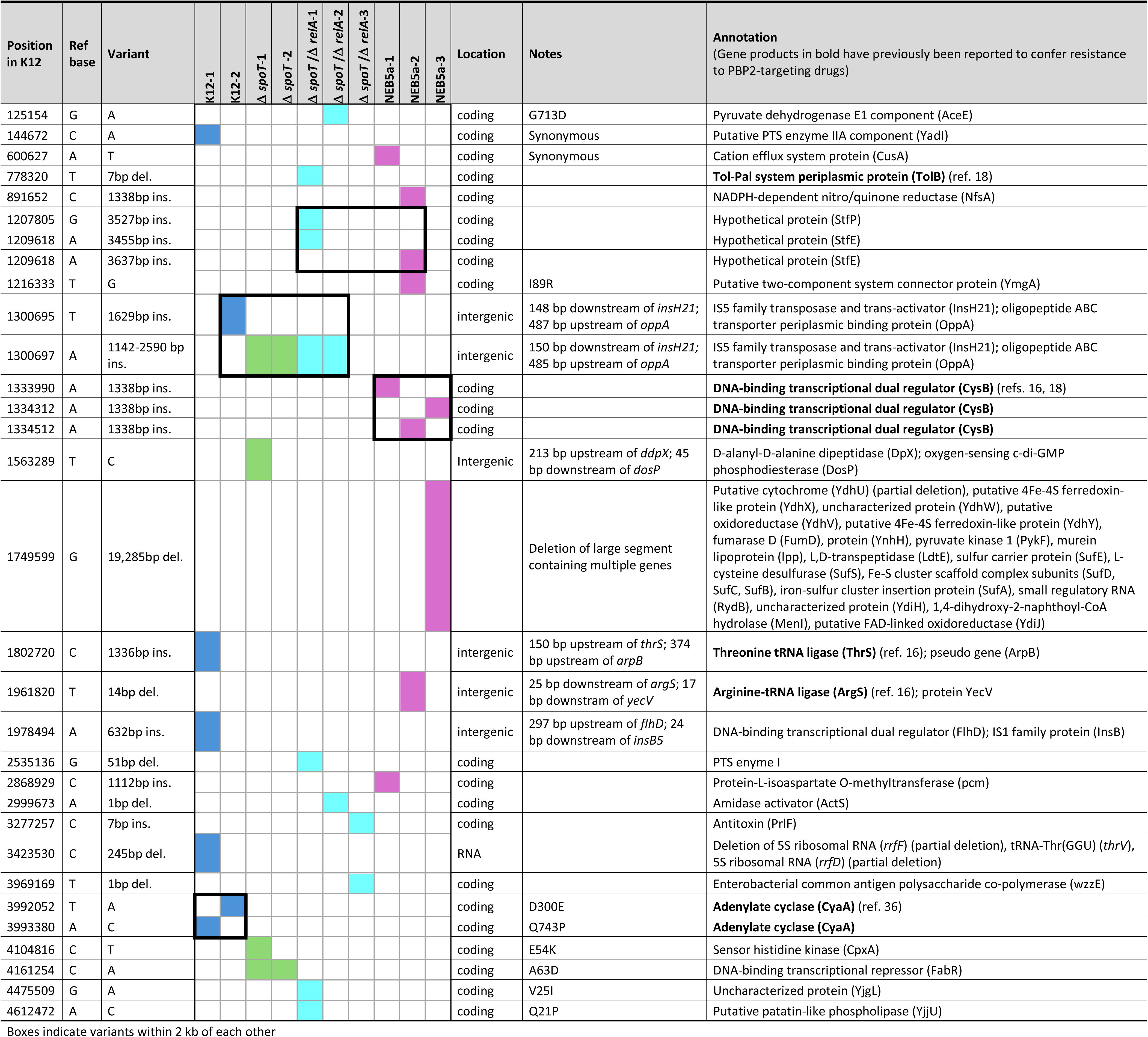
Genetic changes identified in avibactam-resistant strains.

The complexity and variety of AVI resistance-conferring genomic mutations prompted consideration of whether epigenetic changes, in the form of differential methylation, could also be playing a role in resistance. We thus used long-read sequencing technology to quantify methylation of sites throughout the genomes of the strains that had undergone whole-genome sequencing. 5-methylcytosine (5mc) and N6-deoxyadenosine (6ma) methylation was predicted from Oxford Nanopore sequencing data. In total, sites of differential methylation (see Methods for criteria used to identify these sites) occurred in 448 different genes and 39 intergenic regions in the case of 5mc methylation, but only 11 genes and 9 intergenic regions for 6ma methylation (Tables S3, S4). Sites of differential 6ma methylation were unevenly distributed, with 10 of 35 sites of differential methylation occurring in the putative outer membrane porin gene *nmpC* or the adjacent intergenic region; in all cases, these represented a decrease in methylation in AVI-resistant mutants derived from NEB 5 alpha. Interestingly, this gene is located near *rusA*, which was the most represented gene in 5mc methylation differences (see below). Overall, NEB 5 alpha was greatly overrepresented in 6ma methylation, with 75 sites of differential methylation occurring across the 3 NEB 5 alpha mutants and only 16 in all other mutants combined.

Sites of 5mc differential methylation were spread more evenly across strain backgrounds, with many sites occurring in multiple strain backgrounds (Table S4). The gene with the greatest number of 5mc differences was *rusA*, which encodes an endonuclease that resolves Holliday Junction intermediates created during DNA repair by homologous recombination (40). In 2 of the NEB 5 alpha mutants, *rusA* 5mc methylation was decreased at five different sites, while in all three Δ*spoT*/Δ*relA* mutants, methylation in this gene was increased at a separate site. *RuvC*, another such “resolvase” (41), was also differentially methylated in several samples, with decreased methylation in two NEB 5 alpha mutants and one K12 mutant and increased methylation in two Δ*spoT*/Δ*relA* mutants. While these data do not provide information on whether methylation at these sites resulted in increased or decreased gene expression, the predominance of genes involved in DNA repair is notable. Many of the genes that have previously been implicated in resistance to PBP2-targeting drugs were also differentially methylated, including several tRNA ligases (*alaS, asnS, proS, serS, thrS*), as well as *cyaA* (adenylate cyclase), *cysE*, and *ubiX* (16), and *arcA*, *cydA*, and *tolB* (18).

### Avibactam resistance confers a modest *in vitro* fitness cost

In a growth rate assay performed to assess for potential fitness costs of AVI resistance, 2 AVI-resistant mutant derivatives of *E. coli* K12 showed increased lag time relative to the parent strain, but did not have a statistically significant decrease in growth rate (Figure 7). Resistance to PBP2-targeting drugs involves compensatory mechanisms that allow survival and replication in the presence of PBP2 inhibition, but the abnormal morphology of cells grown in the presence of these drugs (Figure 3b) suggests the possibility of a further fitness cost during drug exposure. To evaluate this possibility, we also tested growth fitness of the two AVI-resistant isolates in the presence of a sub-MIC AVI concentration (128 μg/mL). Both strains showed an increase in lag time when grown with AVI, but only one of the strains (mutant #2) demonstrated a significantly decreased growth rate. Interestingly, both of these strains have mutations in the gene encoding adenylate cyclase, but mutant #1 also has an insertion sequence 145 bp upstream of *thrS* (threonine-tRNA ligase), potentially in the promoter region (Table 3). These results suggest that different collections of AVI-resistance mutations may confer differential fitness costs.

**Figure 7.**
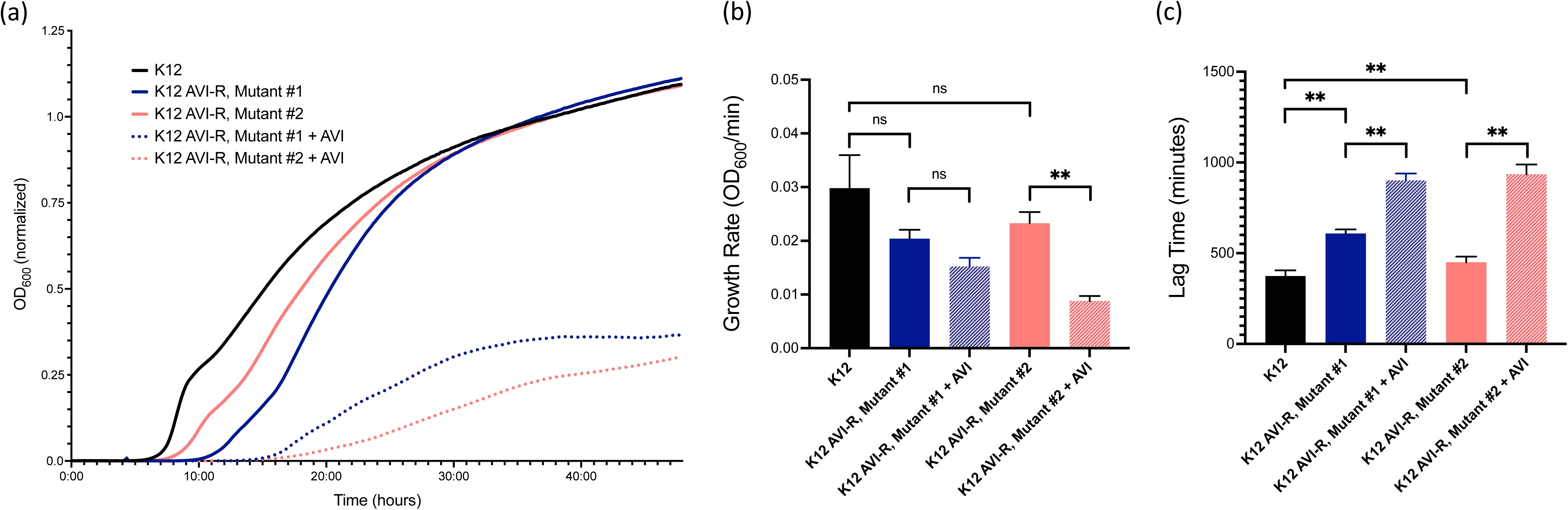
Growth fitness assay demonstrating growth over time of *E. coli* K12 and avibactam-resistant (AVI-R) mutant derivative strains grown with and without AVI 128 μg/mL. (**a**) Growth curves. Data represent mean of three biological replicates. Readings are normalized to media-only wells. (**b-c**) Growth rates and lag time. Measurements in Figs 7b-c were calculated using GrowthRates 6.2.1 (Bellingham Research Institute). Mean values with standard deviation are shown. **, p <0.01; ns, nonsignificant via paired two-tailed t-test.

### Avibactam appears to have greater *in vivo* efficacy in an immunocompetent mouse model

In the neutropenic thigh infection model, mice infected with *K. pneumoniae* FDA-CDC 0636 and treated with 250 mg/kg AVI every 8 hours for 3 doses had a bacterial burden at 24 hours that was 0.5 log_10_ CFU/thigh lower than in mice treated with saline (Mann-Whitney U = 0; *p* = 0.029) (Figure 8). To evaluate the possible effect of an intact innate immune response on AVI activity, AVI was also evaluated in a non-neutropenic thigh infection model. In immunocompetent mice, AVI had greater activity, with treated mice showing a bacterial burden 1.7 log_10_ CFU/thigh lower than saline-treated mice (Mann-Whitney U = 0.5; *p* = 0.016) (Figure 8).

**Figure 8.**
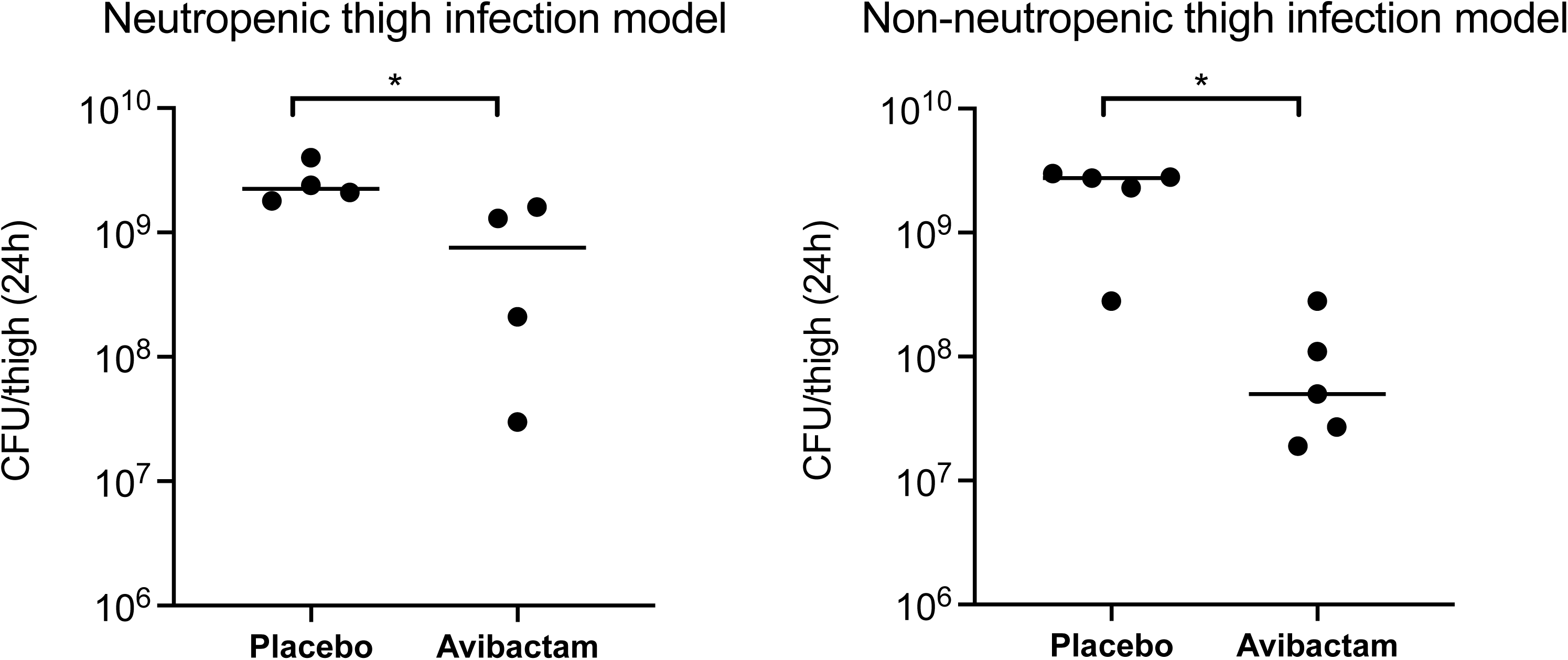
Colony counts of *K. pneumoniae* FDA-CDC 0636 (“Nevada” strain) at 24 hours in mice treated with placebo or avibactam. *, p <0.05 via Mann-Whitney U test.

## DISCUSSION

The development of β-lactam/β-lactamase inhibitor combinations in which the β-lactamase inhibitor is a DBO compound possessing potent direct antimicrobial activity (e.g. zidebactam, nacubactam) suggests the tantalizing possibility of a novel therapeutic approach for infections caused by MBL-producing gram-negative bacteria, for which there are few existing treatment options. Early case reports of compassionate use of cefepime-zidebactam for treatment of MBL-producing *P. aeruginosa* infections have been encouraging (13, 14). However, the activity of these combinations against MBL-producing strains relies on inhibition of PBP2 by the DBO, as DBOs do not inhibit MBLs and therefore cannot protect their partner β-lactam drug against MBL-mediated hydrolysis (42). Even the β-lactam “enhancer” effect, in which DBO/β-lactam combinations retain activity against some strains that are resistant to both the β-lactam and the DBO alone, is believed to result from synergistic activity between residual low-level inhibition of PBP2 by the DBO and inhibition of other PBPs by the partner β-lactam, thus continuing to rely on PBP2 inhibition as a component of intrinsic activity (17, 42, 43).

There is, however, reason for concern that reliance on PBP2 inhibition for antibacterial activity is a tenuous strategy. The only currently clinically available drug with selective activity against PBP2 is mecillinam, a β-lactam antibiotic for which gram-negative bacteria have such a high mutation frequency that it is only used to treat bladder infections, where its concentration at extremely high levels in urine limits resistance (44). However, urine mecillinam concentrations are at least 10 times higher than zidebactam concentrations in the bloodstream at doses used in human trials (45), such that resistance may be a greater threat when novel DBOs are used to treat invasive infections. We found that resistance to AVI emerged readily upon exposure to the drug at concentrations as much as 16 times the MIC. This phenomenon first became apparent in time-kill studies, where bacterial regrowth occurred reliably by 24 hours (Figures 2 and 6). (The explanation for the discrepancy between inhibition of growth in the MIC assay and failure of inhibition in the time-kill assay can be understood by considering the baseline AVI resistance frequency of 1.97 ×10^−6^ – 8.15 ×10^−5^ among the strains we tested. The starting bacterial inoculum in time-kill studies is ~10^8^ cells (10^6^ CFU/mL x 10 mL volume), whereas in the 384-well plate MIC method it is 2.5×10^4^ (5×10^5^ CFU/mL x 0.05 mL volume), thus at least one AVI-resistant cell is almost certain to occur in a time-kill study, but not in an MIC assay.) AVI resistance persisted over more than two weeks of serial subcultures in the absence of antibiotic pressure, demonstrating that cells had developed true heritable resistance, and there was no evidence of tolerance when this was tested directly.

As expected, resistance to AVI conferred resistance to other exclusively PBP2-targeting drugs, but not to β-lactam antibiotics with different or additional PBP targets (Table 2). The morphological appearance of AVI-resistant cells grown in the presence of AVI was consistent with the enlarged, round forms seen in cells resistant to other PBP2-targeting drugs, including mecillinam (23, 46) and nacubactam (18) (Figure 3b). The fact that AVI-resistant and AVI-susceptible cells take on the same distorted appearance in the presence of AVI is reflective of the remarkable approach taken by bacteria in developing resistance to PBP2-inhibiting drugs. In the great majority of cases, resistance to PBP2-targeting drugs in *Enterobacterales* occurs via the emergence of one or more of a large number of apparently compensatory mutations, which allow bacteria to survive and replicate in the presence of PBP2 inhibition, rather than preventing PBP2 inhibition outright (16). This large mutational target is believed to confer resistance to PBP2 inhibitors by upregulation of the stringent response (47). Activation of the *ftsAQZ* operon and resultant ability of bacterial cells to survive and replicate as the enlarged, rounded forms generated by PBP2 inhibition has been proposed as the final step leading from stringent response activation to resistance to PBP2 inhibition (18), although the process, and the role of the many different mutations identified in isolates resistant to PBP2 inhibition, remains incompletely understood. In two AVI-resistant mutants derived from *E. coli* K12, we found non-synonymous point mutations in *cyaA*, the gene encoding adenylate cyclase, which has previously been described as a cause of mecillinam resistance (36, 48), potentially mediated by effects on lipopolysaccharide synthesis (48). One of the strains also had an insertion sequence in the *thrS* gene encoding threonine-tRNA ligase. Mutations in tRNA synthetases are among the most common genetic changes identified in bacteria resistant to PBP2 inhibiting drugs (16, 49) and are believed to simulate amino acid starvation, resulting in a stimulus for upregulation of the stringent response (18). To evaluate emergence of AVI resistance in a different *E. coli* K12 strain background, we sequenced 3 AVI-resistant mutants of NEB 5-alpha, a derivative of DH5α (50). Interestingly, all three of these isolates had an insertion sequence within *cysB*. Inactivation of CysB, a positive regulator of cysteine biosynthesis, is identified frequently in clinical mecillinam-resistant *E. coli* isolates, but rarely in standard laboratory-selected mutants, potentially because of increased growth fitness in urine (16). CysB inactivation is believed to confer mecillinam resistance through a pathway that is independent of the stringent response but has similar downstream effects, ultimately rendering PBP2 inessential (51). Although it is not clear why this mutation would be preferentially selected in the NEB 5-alpha strain background, this finding suggests that differences in strain characteristics may have important impacts on the type of mutations that emerge to PBP2-targeting drugs and, in turn, on the fitness and potential clinical importance of these isolates. Because sequencing for each parent strain was performed on isolated colonies generated from a single starting culture, the detection of multiple strains with the same mutation could reflect the emergence of resistant mutants at one or more potential time points: (1) pre-existing mutations present at a low level within the parent strain population and selected under AVI pressure, (2) early emergence of a mutant clone giving rise to multiple colonies or (3) convergent evolution of the same mutation in multiple different cells. Future experiments, potentially using methods in development such as single-cell whole genome sequencing, may be able to clarify the time course of emergence of resistance.

The fact that mutations causing upregulation of the stringent response predominate in laboratory-selected strains resistant to PBP2 inhibition raised the question of whether, and how, strains deficient in key genes in the stringent response pathway might develop AVI resistance. Surprisingly, we saw no significant decrease in mutation frequency in strains lacking *spoT*, *relA*, or both (Figure 5). However, the genes altered in AVI-R mutants of these strains were, with one exception (a 7-base pair deletion in *tolB* in one of the Δ*spoT*/Δ*relA* double mutants (18)), genes that have not previously been described in bacteria resistant to PBP2-targeting drugs and, as expected, are not part of the stringent response pathway. TolB is a component of an alternate path that may result in activation of the *ftsQAZ* operon (18), but to our knowledge, the other genes in which we identified mutations are not known to participate in this process. Our analysis of differences in methylation between AVI-resistant mutants and their parent strains highlighted several of the same genes in which mutations are frequently found in PBP2 inhibitor resistance, as well as several genes not previously implicated in resistance to these drugs. Our cumulative data underscore the strikingly complex and multifarious mechanisms by which bacteria develop resistance to PBP2-targeting drugs, although the reason for this unusual approach to resistance remains opaque.

Our study has certain limitations. We have not yet induced mutations in the novel genes we identified to confirm that their inactivation confers AVI resistance; this will be an important step in future work. In addition, our analysis of methylation data does not provide information on whether genes with increases in methylation are silenced, and more detailed investigation of methylation patterns and gene expression levels in the future will help to elucidate the role of methylation in resistance to PBP2-targeting drugs.

One important consideration with any *in vitro* study of antibiotic resistance is the extent to which resistance phenotypes identified in laboratory settings will translate to clinical resistance and treatment failure. We did observe a fitness cost, in terms of lag time but not growth rate, in AVI-resistant isolates (Figure 7), which might suggest that AVI-resistant isolates would be less likely to survive in the more exacting host environment. The fact that AVI appeared more active in an immunocompetent mouse model compared to an otherwise identical neutropenic model (Figure 8) provides support for the idea that host responses may reduce the likelihood of emergence of resistance. Indeed, Ulloa et al. have noted that components of the innate immune system exert synergistic activity with AVI against MBL-producing *K. pneumoniae* (52). However, the large mutational target leading to AVI resistance suggests that bacteria exposed to DBOs *in vivo* may preferentially select options from the “menu” of mutations that allow for improved survival in the presence of AVI (Figure 7) and under various selective pressures exerted by the host. Future studies investigating the ability of AVI-resistant mutants to cause infection in preclinical models may improve our ability to predict the effect of resistance *in vivo*.

The rapidity with which resistance to AVI emerges, as well as the complex and incompletely understood collection of mutations capable of conferring such resistance, should be considered a warning sign as novel β-lactam/DBO combinations become available for clinical use. These new agents offer an enticingly broad spectrum of activity against MDR gram-negative bacteria, but the reliance of this activity on direct inhibition of PBP2 by DBOs raises concerns about the durability of the new drugs’ activity. Future studies characterizing rates and mechanisms of resistance to DBO-containing β-lactam/β-lactamase inhibitor combinations, studies in preclinical models, and close observation of clinical treatment-emergent resistance will be essential in preserving the activity of these drugs in the ongoing fight against MDR gram-negative pathogens.

## MATERIALS AND METHODS

### Bacterial Strains

Bacteria were obtained from the following sources (Table S1 and S2): the U.S. FDA-CDC Antimicrobial Resistance Isolate Bank (48 isolates), the Antibiotic Resistance Leadership Group Laboratory Center Virtual Repository (1 isolate), the carbapenem-resistant *Enterobacteriaceae* genome initiative at the Broad Institute in Cambridge, MA (21 isolates) (53, 54), New England BioLabs (1 isolate), and the Coli Genetic Stock Center (4 isolates). *Escherichia coli* ATCC 25922, *Staphylococcus aureus* ATCC 29213, *Klebsiella pneumoniae* ATCC 700603, *K. pneumoniae* ATCC 13883, *K. pneumoniae* ATCC BAA-1705 and *Pseudomonas aeruginosa* ATCC 27853 were obtained from the American Type Culture Collection (Manassas, VA). All strains were colony purified, minimally passaged, and stored at −80°C in tryptic soy broth (BD Diagnostics, Franklin Lakes, NJ) with 50% glycerol (Sigma-Aldrich, St. Louis, MO) prior to use in this study.

### Antimicrobial Agents

Ceftazidime and cefepime were obtained from Chem Impex International (Wood Dale, IL). Avibactam was obtained from MedChemExpress (Monmouth Junction, NJ). Clavulanate was obtained from Sigma-Aldrich (St. Louis, MO). Amoxicillin was obtained from Alfa Aesar (Tewksbury, MA). Mecillinam (amdinocillin) was obtained from Research Products International (Mt. Prospect, IL). Meropenem was obtained from Ark Pharm, (Libertyville, IL). Aztreonam was obtained from MP Biomedicals (Solon, OH). Antimicrobial stock solutions used for the digital dispensing method (DDM) were dissolved in water or in dimethyl sulfoxide (DMSO) according to Clinical and Laboratory Standards Institute (CLSI) recommendations (55); 0.3% polysorbate 20 (P-20; Sigma-Aldrich, St. Louis, MO) was added to the water used for these solutions as required by the HP D300 digital dispenser instrument (HP, Inc., Palo Alto, CA) for proper fluid handling. As recommended by CLSI, anhydrous sodium carbonate at 10% ceftazidime weight was added to the ceftazidime stock solution. The antibiotic stock solutions used for time-kill studies and agar dilution plates were dissolved in water, with the addition of anhydrous sodium carbonate at 10% weight/weight for ceftazidime. All antibiotic stock solutions were QC tested with *E. coli* ATCC 25922, *S. aureus* ATCC 29213, *K. pneumoniae* ATCC 700603, and/or *K. pneumoniae* ATCC BAA-1705 using the D300 dispensing method described below (for stocks to be used for checkerboard arrays) or standard broth microdilution using the direct colony suspension method (56) (for stocks to be used for time-kill experiments) prior to use in synergy studies. Stocks were used only if they produced an MIC result within the accepted QC range according to CLSI guidelines (55). Because the MIC of avibactam for *K. pneumoniae* FDA-CDC 0636 was noted to be consistent at 8 μg/mL, this strain was used as an alternate QC organism to reduce the variables involved in QC of avibactam in combination with ceftazidime. Antimicrobials were stored as aliquots at −20°C and discarded after a single use, except for ceftazidime, which was either prepared fresh the day of use or stored at −80°C.

### Minimal Inhibitory Concentration (MIC) and Checkerboard Array Synergy Testing

Digital dispensing method (DDM) MIC testing was performed using the HP D300 digital dispenser (HP, Inc., Palo Alto, CA, USA) as previously described by our laboratory (57, 58). Bacterial inocula were adjusted to a McFarland reading of 0.5 and diluted 1:300 in cation-adjusted Mueller Hinton broth (CAMHB), resulting in a suspension of ~5×10^5^ CFU/mL, and 50 μL of this suspension was added to wells in flat-bottomed, untreated 384-well polystyrene plates (Greiner Bio-One, Monroe, NC, USA) using a multichannel pipette. Antimicrobial stock solutions were dispensed by the D300 into wells before or immediately after addition of bacterial suspensions. Plates were incubated at 35-37°C in ambient air for 16–20 h. After incubation, bacterial growth was quantified by measurement of OD_600_ using a Tecan Infinite M1000 Pro microplate reader (Tecan, Morrisville, NC, USA). An OD_600_ reading of >0.08 (approximately twice typical background readings in wells containing broth alone) was considered indicative of bacterial growth; this cutoff correlated with inhibition of growth by visual assessment.

### Time-Kill Studies

Antibiotic stocks for time-kill studies were prepared as described above and diluted in 10 mL of CAMHB in 25- by 150-mm glass round-bottom tubes to the desired starting concentrations. The starting inoculum for time-kill studies was prepared by adding 100 μl of a 1.0 McFarland standard suspension of colonies from an overnight plate to each of the tubes. A growth control and a negative (sterility) control tube were prepared in parallel with each experiment. Cultures were incubated on a shaker in ambient air at 35-37°C. Aliquots from the culture were removed at serial time points, and a 10-fold dilution series was prepared in 0.9% sodium chloride. A 10 μl drop from each dilution was transferred to a Mueller-Hinton agar plate (ThermoFisher, Waltham, MA) and incubated overnight in ambient air at 35-37°C (59). For countable drops (drops containing 3 to 30 colonies), the cell density of the sample was calculated; if more than one dilution for a given sample was countable, the cell density of the two dilutions was averaged. If no drops were countable, the counts for consecutive drops above and below the countable range were averaged. The lower limit of quantitation was 300 CFU/mL.

### Persistence of Avibactam Resistance

Following growth in CAMHB with either no drug or 128 μg/mL AVI for 24 hours as in the time-kill method, liquid cultures were transferred to 15 mL conical centrifuge tubes and centrifuged at 5000g for 10 minutes at 24°C. The supernatant from each tube was poured off and cells were resuspended in fresh CAMHB, then diluted to a 0.5 McFarland standard in 0.9% sodium chloride. AVI MIC testing was performed on these suspensions using the DDM described above; in addition, bacteria from each suspension were isolation streaked onto a TSA/5% sheep blood agar plate (ThermoFisher). Following overnight incubation in ambient air at 35°C, isolated colonies from these plates were used to perform AVI MIC testing and for isolation streaking onto new blood agar plates. This procedure was repeated for 15 days, with MIC testing performed at 7 points within this time frame. Three replicates of the entire procedure were performed with each strain.

### Biological Assay to Determine Avibactam Concentration in Media after 24 Hours

Tubes containing bacteria (*K. pneumoniae* FDA-CDC 0636 and *E. coli* ARLG 2829) and AVI at 128 μg/mL were prepared and incubated as in time-kill studies. After 24 hours, 7 mL from each tube was transferred to a 15 mL conical tube and centrifuged at 5000g for 10 minutes at 24°C. Three mL of the resulting supernatant were then removed from each tube and sterilized using a syringe filter. A serial two-fold dilution series was prepared in CAMHB in a 96-well untreated round-bottom plate, with the first well in each row containing undiluted supernatant. An additional row containing a dilution series of AVI stock of known concentration was prepared in parallel. Bacterial suspensions of both strains were then prepared and added to wells at a final concentration of 5×10^5^ CFU/mL. The plates were incubated overnight. The lowest concentration in the supernatant dilution series in which growth was inhibited was assumed to contain approximately the same concentration of drug as the MIC in the known-concentration row, and this data was used to calculate the concentration of drug in the undiluted well.

### Mutation Frequency Analysis

Agar dilution plates were prepared by adding one part of a 10X antibiotic concentration to nine parts molten 1.5% Bacto agar (Becton, Dickinson and Company, Sparks, MD, USA) containing CAMHB. Bacterial strains were streaked onto TSA/5% sheep blood agar plates and incubated overnight in ambient air at 37°C. A single colony of each strain was added to 10 mL of CAMHB in a 25- by 150-mm glass round bottom tube and incubated overnight with shaking in ambient air at 37°C. A 150 µL aliquot was then removed from each culture tube and used to prepare a 1:10 dilution in 0.9% NaCl. A 10 µL spot from each dilution was placed on the agar plates with a multichannel pipette, left to dry, and incubated overnight or until visible colonies were apparent. Colonies were counted as described above and the ratio of CFU at each antibiotic concentration to CFU in the non-antibiotic-containing plate was calculated.

### Tolerance Assay

Bacterial tolerance was assessed using the TDtest described by Gefen et al (30). AVI-containing disks were prepared by applying 10 µL of an 8 mg/mL stock solution of AVI (total quantity 80 µg; determined through initial assays to produce a zone of ~20 mm with *E. coli* K12) to a diffusion disk (BD Diagnostics, Franklin Lakes, NJ). Glucose disks were prepared by applying 5 µL of a filter-sterilized solution of 40% glucose to a disk. A 0.5 McFarland bacterial inoculum was prepared and spread onto Mueller-Hinton agar plates using Dacron swabs to create a bacterial lawn. An AVI-containing disk was placed onto the plate, which was incubated at 37 degrees Celsius overnight. The avibactam disk was then removed and replaced with a glucose-containing disk and again incubated overnight. Tolerance was assessed by visually inspecting the plates for bacterial growth within the zone of inhibition after the second night of incubation.

### Generation of AVI-resistant Mutants for Growth Rate Assay and Sequencing

Parent strains were isolation streaked onto TSA/5% sheep blood agar plates and incubated overnight at 35-37°C. The following day, liquid cultures were set up from a single colony from each of the strains in 5 mL of CAMHB and incubated overnight with shaking. Cultures were then diluted to a 1.0 McFarland standard (~3×10^8^ CFU/mL) in saline, spread with beads on Mueller Hinton agar plates containing AVI at 128 μg/mL, and incubated overnight. The next day, 3 individual colonies from each plate were separately isolation streaked onto plates containing AVI at 128 μg/mL. When colonies of different morphologies (typically different sizes) appeared on the original plate, colonies of different sizes were selected to increase the likelihood of genetic diversity. Bacterial growth from these plates was then frozen in glycerol stocks at −80°C.

### Growth Rate Assay

*E. coli* K12 and 2 AVI-resistant mutant derivatives were streaked onto agar plates; for the AVI-resistant strains the agar contained AVI at 128 μg/mL. Plates were incubated overnight, and the following day bacterial inocula of approximately 1,000 CFU/mL were prepared in CAMHB from growth on the plates, and 100 μL was added to wells in a 96-well plate, for a total of approximately 100 cells per well. AVI at 128 μg/mL was added to select wells using the DDM. Twelve wells were prepared for each condition (*E. coli* K12 without antibiotic and 2 resistant mutants with and without AVI); the remaining wells contained CAMHB alone. The plate was incubated at 37°C for 48 hours in the Tecan Infinite M1000 Pro microplate reader, with automatic OD_600_ readings obtained every 10 minutes following 10 seconds of orbital shaking. Three biological replicates of the assay were performed. The data was analyzed using GrowthRates 6.2.1 (Bellingham Research Institute) (60).

### Isolation of Genomic DNA

AVI-resistant mutants were generated from *E. coli* K12, two K12-derived strains with mutations in stringent response pathway genes (Δ*spoT*, Δ*spoT*/Δ*relA*), and NEB® 5-alpha, a DH5α derivative electrocompetent cloning strain in which the SOS *recA* gene is inactivated (50). To isolate genomic DNA, AVI-resistant strains were streaked from frozen stocks onto Mueller Hinton agar plates containing AVI at 128 μg/mL and parent (AVI-susceptible) strains were streaked onto TSA/5% sheep blood agar plates and incubated overnight. Colonies from each plate were added to glass tubes containing 13 mL of CAMHB either with (AVI-R strains) or without (parent strains) 128 μg/mL AVI and incubated overnight with shaking. The following day, DNA extraction was performed using QIAGEN Genomic-tip 100/G gravity flow anion-exchange tips (QIAGEN, Germantown, MD) and QIAGEN buffers according to kit instructions. In brief, cells were pelleted by spinning culture in 15 mL conical tubes at 3000-5000g for 5-10 minutes at 21°C. Supernatant was then discarded and cells were resuspended in 3.5 mL Buffer B1 with 200 μg/mL RNAse A (Monarch, New England Biolabs, Ipswich, MA) and vortexed thoroughly. Eighty microliters of 100 mg/mL lysozyme (ThermoFisher) and 100 μL of 20 mg/mL proteinase K (ThermoFisher) were added to each sample, samples were incubated at 37°C for 30 minutes, then 1.2 mL of Buffer B2 was added and samples were mixed and incubated at 50°C for 30 minutes. At this point, each G/100 tip was equilibrated with 4 mL of QBT and allowed to fully empty, then the clarified lysate was added to each column and left overnight to allow binding to the resin and flow-through of the remainder of cell constituents. The columns were then washed twice with 7.5 mL QC buffer. The DNA was eluted in 5 mL of pre-warmed QF buffer and precipitated by adding 3.5 mL room temperature isopropanol and inverting. The DNA was then spooled on a metal inoculating loop, transferred to a 1.5 mL tube containing 200 μL TE, and resuspended overnight at 4°C. Initial evaluation of DNA purity and quantity was performed using the Thermo Scientific™ NanoDrop 2000. Sufficient DNA for sequencing was extracted from two mutants each of *E. coli* K12 and the Δ*spoT* strain and three mutants each of the Δ*spoT*/Δ*relA* strain and NEB® 5-alpha.

### Genome Sequencing, Assembly, and Analysis

Illumina libraries were constructed using the Illumina Nextera XT protocol and sequenced using the Illumina NovaSeq 6000 platform to a depth of approximately 1.5 gigabases per sample. Illumina reads for parental and mutant strains were processed using Trimmomatic version 0.39 (61), then aligned against the *E. coli* K12 reference genome (NCBI Genbank accession GCA_000005845.2) using BWA Mem version 0.7.17 (62). Single nucleotide polymorphisms (SNPs) and structural variation, like insertions and deletions, were called using Pilon version 1.23 (35). SNPs identified as “Passing” by Pilon were used in downstream analyses except when SNPs were common to parental and mutant strains; regions with variable length indels in both the parental and descendent strains, and variants identified by Pilon as duplications were also excluded.

Oxford Nanopore Technologies (ONT) long-read sequencing libraries were constructed using the Oxford Nanopore kit SQK-LSK109. Samples were barcoded and run in batches of 12 on a GridION machine (Oxford Nanopore Technologies Ltd, Science Park, UK). Initial processing of reads was performed as previously described (54). To call methylation states on the ONT data, we ran Guppy version 6.1.7 (https://community.nanoporetech.com) with the Rerio (https://github.com/nanoporetech/rerio) model res_dna_r941_min_modbases-all-context_v001.cfg, outputting 5mC and 6mA methylation information aligned to the *E. coli* K12 reference genome. We used modbam2bed (https://github.com/epi2me-labs/modbam2bed) to further process the output from Guppy. Sites in which methylation patterns differed in AVI-resistant strains compared to parent strains were identified using the following criteria: (1) the parent strain had coverage of ≥3 reads at the position (average was ~85 reads/position), (2) the difference in percent methylation between parent and mutant strain was at least 50%, and (3) at least two different mutant samples met these criteria. Methylation differences in intergenic regions were considered to have potentially affected both adjacent genes.

Gene annotations were those provided in the NCBI sequence of *E. coli* K12 substr. MG1655 (NCBI Genbank identifier U00096).

### Murine Thigh Infection Models

#### Neutropenic Thigh Infection Model

Twelve female CD-1 (ICR) mice (Charles River Laboratories, Cambridge, MA) weighing 25-30 grams were treated with cyclophosphamide (European Pharmacopoeia Reference Standard) by intraperitoneal (IP) injection (150 mg/kg on day −4 and 100 mg/kg on day −1) to induce neutropenia (63, 64). On day −3, mice were treated with 5 mg/kg uranyl nitrate to cause renal impairment simulating human drug clearance (65). On day 0, a suspension of approximately 1×10^7^ CFU/mL of *K. pneumoniae* FDA-CDC 0636 was prepared in sterile endotoxin-free 0.9% sodium chloride (Teknova, Hollister, CA). Mice were anesthetized with isoflurane and injected in the right thigh with 100 μL of the bacterial suspension (total 1×10^6^ CFU/thigh). Four of the mice were then sacrificed by CO_2_ inhalation for baseline colony enumeration. (In one mouse, the thigh injection was inadvertently performed subcutaneously; data from this mouse was not used in subsequent calculations). Following sacrifice, the right thigh was dissected, suspended in 1 mL sterile 0.9% sodium chloride, and emulsified in a tissue grinder. A sample of the liquid homogenate was then removed for serial 10-fold dilutions in 0.9% sodium chloride, plating, and colony enumeration using the drop method described above. Approximately three hours after the time of bacterial thigh infection, the remaining mice were injected subcutaneously with either 250 mg/kg of AVI dissolved in 0.9% sodium chloride (4 mice) or an equivalent volume of sodium chloride alone (4 mice); doses were repeated twice after this at 8-hour intervals for a total of 3 doses. AVI dosing was selected based on the highest AVI dosing reported in the literature in mouse thigh models of AVI in combination with ceftazidime (66). At 24 hours after the first dose, mice were euthanized and thighs removed for colony enumeration as described above. (One mouse in the AVI treatment group developed lethargy and apparent seizure activity shortly after the third AVI injection, at which time it was sacrificed; thigh dissection and plating were performed immediately following sacrifice).

#### Non-neutropenic Thigh Infection Model

To determine the bacterial burden required to establish a thigh infection with *K. pneumoniae* FDA-CDC 0636 in a non-neutropenic mouse, 2 mice each were injected with three different bacterial inocula (1×10^6^, 1×10^7^ and 1×10^8^ CFU/thigh) as described above. After 24 hours, mice were euthanized and thighs removed for colony enumeration as described above; only the highest inoculum resulted in an increase in CFU/thigh over the 24 hour period (increase in CFU/thigh of 1.3 log_10_ vs decrease of ~1.8 log_10_ CFU/thigh with the two lower inocula). The treatment experiment was performed as for the neutropenic thigh infection model, except that pre-treatment with cyclophosphamide was omitted and the 10^8^ CFU/thigh inoculum was used for infection. There were five mice each in the baseline, AVI treatment, and saline treatment groups.

All mouse experiments were performed under an institutional animal care and use committee (IACUC)-approved protocol.

### Data Analysis

Statistical analysis was performed using GraphPad Prism 10 software. Growth curve data analysis was performed using GrowthRates 6.2.1 (Bellingham Research Institute) (60).

### Data Availability

Both Illumina and Oxford Nanopore sequencing reads were deposited at the Sequence Read Archive under Bioproject PRJNA1140646 (https://www.ncbi.nlm.nih.gov/bioproject/?term=PRJNA1140646).

## ACKNOWLEDGEMENTS

We would like to thank Terrence Shea for helpful discussions. This project has been funded in part with Federal funds from the National Institute of Allergy and Infectious Diseases, National Institutes of Health, Department of Health and Human Services, under Grant Numbers K08AI132716 and R01AI178875 to Thea Brennan-Krohn and U19AI110818 to the Broad Institute.

## References

1. Bush K, Bradford PA. 2016. β-Lactams and β-Lactamase Inhibitors: An Overview. Cold Spring Harb Perspect Med 6:a025247.

2. Nordmann P, Naas T, Poirel L. 2011. Global spread of Carbapenemase-producing Enterobacteriaceae. Emerg Infect Dis 17:1791–1798.

3. Drawz SM, Bonomo RA. 2010. Three decades of beta-lactamase inhibitors. Clin Microbiol Rev 23:160–201.

4. Ehmann DE, Jahić H, Ross PL, Gu R-F, Hu J, Kern G, Walkup GK, Fisher SL. 2012. Avibactam is a covalent, reversible, non-β-lactam β-lactamase inhibitor. Proc Natl Acad Sci U S A 109:11663–11668.

5. Zhanel GG, Lawson CD, Adam H, Schweizer F, Zelenitsky S, Lagacé-Wiens PRS, Denisuik A, Rubinstein E, Gin AS, Hoban DJ, Lynch JP, Karlowsky JA. 2013. Ceftazidime-avibactam: a novel cephalosporin/β-lactamase inhibitor combination. Drugs 73:159–177.

6. Asli A, Brouillette E, Krause KM, Nichols WW, Malouin F. 2016. Distinctive Binding of Avibactam to Penicillin-Binding Proteins of Gram-Negative and Gram-Positive Bacteria. Antimicrob Agents Chemother 60:752–756.

7. Sauvage E, Kerff F, Terrak M, Ayala JA, Charlier P. 2008. The penicillin-binding proteins: structure and role in peptidoglycan biosynthesis. FEMS Microbiol Rev 32:234–258.

8. Mushtaq S, Vickers A, Woodford N, Haldimann A, Livermore DM. 2019. Activity of nacubactam (RG6080/OP0595) combinations against MBL-producing Enterobacteriaceae. J Antimicrob Chemother 74:953–960.

9. Peilleron L, Cariou K. 2020. Synthetic approaches towards avibactam and other diazabicyclooctane β-lactamase inhibitors. Org Biomol Chem 18:830–844.

10. Giri P, Patel H, Srinivas NR. 2019. Review of Clinical Pharmacokinetics of Avibactam, A Newly Approved non-β lactam β-lactamase Inhibitor Drug, In Combination Use With Ceftazidime. Drug Res 69:245–255.

11. Livermore DM, Mushtaq S, Warner M, Vickers A, Woodford N. 2017. In vitro activity of cefepime/zidebactam (WCK 5222) against Gram-negative bacteria. J Antimicrob Chemother 72:1373–1385.

12. Study of Cefepime-zidebactam (FEP-ZID) in Complicated Urinary Tract Infection (cUTI) or Acute Pyelonephritis (AP). https://clinicaltrials.gov/study/NCT04979806.

13. Tirlangi PK, Wanve BS, Dubbudu RR, Yadav BS, Kumar LS, Gupta A, Sree RA, Challa HPR, Reddy PN. 2023. Successful Use of Cefepime-Zidebactam (WCK 5222) as a Salvage Therapy for the Treatment of Disseminated Extensively Drug-Resistant New Delhi Metallo-β-Lactamase-Producing Pseudomonas aeruginosa Infection in an Adult Patient with Acute T-Cell Leukemia. Antimicrob Agents Chemother 67:e0050023.

14. Dubey D, Roy M, Shah TH, Bano N, Kulshrestha V, Mitra S, Sangwan P, Dubey M, Imran A, Jain B, Velmurugan A, Bakthavatchalam YD, Veeraraghavan B. 2023. Compassionate use of a novel β-lactam enhancer-based investigational antibiotic cefepime/zidebactam (WCK 5222) for the treatment of extensively-drug-resistant NDM-expressing Pseudomonas aeruginosa infection in an intra-abdominal infection-induced sepsis patient: a case report. Ann Clin Microbiol Antimicrob 22:55.

15. Morinaka A, Tsutsumi Y, Yamada M, Suzuki K, Watanabe T, Abe T, Furuuchi T, Inamura S, Sakamaki Y, Mitsuhashi N, Ida T, Livermore DM. 2015. OP0595, a new diazabicyclooctane: mode of action as a serine β-lactamase inhibitor, antibiotic and β-lactam “enhancer.” J Antimicrob Chemother 70:2779–2786.

16. Thulin E, Sundqvist M, Andersson DI. 2015. Amdinocillin (Mecillinam) resistance mutations in clinical isolates and laboratory-selected mutants of Escherichia coli. Antimicrob Agents Chemother 59:1718–1727.

17. Livermore DM, Warner M, Mushtaq S, Woodford N. 2016. Interactions of OP0595, a Novel Triple-Action Diazabicyclooctane, with β-Lactams against OP0595-Resistant Enterobacteriaceae Mutants. Antimicrob Agents Chemother 60:554–560.

18. Doumith M, Mushtaq S, Livermore DM, Woodford N. 2016. New insights into the regulatory pathways associated with the activation of the stringent response in bacterial resistance to the PBP2-targeted antibiotics, mecillinam and OP0595/RG6080. J Antimicrob Chemother 71:2810–2814.

19. de Man TJB, Lutgring JD, Lonsway DR, Anderson KF, Kiehlbauch JA, Chen L, Walters MS, Sjölund-Karlsson M, Rasheed JK, Kallen A, Halpin AL. 2018. Genomic Analysis of a Pan-Resistant Isolate of Klebsiella pneumoniae, United States 2016. mBio 9:e00440–18.

20. Chen L, Todd R, Kiehlbauch J, Walters M, Kallen A. 2017. Notes from the Field: Pan-Resistant New Delhi Metallo-Beta-Lactamase-Producing Klebsiella pneumoniae - Washoe County, Nevada, 2016. MMWR Morb Mortal Wkly Rep 66:33.

21. Mediavilla JR, Patrawalla A, Chen L, Chavda KD, Mathema B, Vinnard C, Dever LL, Kreiswirth BN. 2016. Colistin- and Carbapenem-Resistant Escherichia coli Harboring mcr-1 and blaNDM-5, Causing a Complicated Urinary Tract Infection in a Patient from the United States. mBio 7:e01191–16.

22. Sutaria DS, Moya B, Green KB, Kim TH, Tao X, Jiao Y, Louie A, Drusano GL, Bulitta JB. 2018. First Penicillin-Binding Protein Occupancy Patterns of β-Lactams and β-Lactamase Inhibitors in Klebsiella pneumoniae. Antimicrob Agents Chemother 62:e00282–18.

23. Gutmann L, Vincent S, Billot-Klein D, Acar JF, Mrèna E, Williamson R. 1986. Involvement of penicillin-binding protein 2 with other penicillin-binding proteins in lysis of Escherichia coli by some beta-lactam antibiotics alone and in synergistic lytic effect of amdinocillin (mecillinam). Antimicrob Agents Chemother 30:906–912.

24. Barbour AG, Mayer LW, Spratt BG. 1981. Mecillinam resistance in Escherichia coli: dissociation of growth inhibition and morphologic change. J Infect Dis 143:114–121.

25. Fass RJ. 1980. Activity of mecillinam alone and in combination with other beta-lactam antibiotics. Antimicrob Agents Chemother 18:906–912.

26. Spratt BG. 1977. Properties of the Penicillin-Binding Proteins of *Escherichia coli* K12. Eur J Biochem 72:341–352.

27. Papp-Wallace KM, Nguyen NQ, Jacobs MR, Bethel CR, Barnes MD, Kumar V, Bajaksouzian S, Rudin SD, Rather PN, Bhavsar S, Ravikumar T, Deshpande PK, Patil V, Yeole R, Bhagwat SS, Patel MV, Van Den Akker F, Bonomo RA. 2018. Strategic Approaches to Overcome Resistance against Gram-Negative Pathogens Using β-Lactamase Inhibitors and β-Lactam Enhancers: Activity of Three Novel Diazabicyclooctanes WCK 5153, Zidebactam (WCK 5107), and WCK 4234. J Med Chem 61:4067–4086.

28. Noguchi H, Matsuhashi M, Mitsuhashi S. 1979. Comparative Studies of Penicillin-Binding Proteins in Pseudomonas aeruginosa and Escherichia coli. Eur J Biochem 100:41–49.

29. Davies TA, Page MGP, Shang W, Andrew T, Kania M, Bush K. 2007. Binding of ceftobiprole and comparators to the penicillin-binding proteins of Escherichia coli, Pseudomonas aeruginosa, Staphylococcus aureus, and Streptococcus pneumoniae. Antimicrob Agents Chemother 51:2621–2624.

30. Gefen O, Chekol B, Strahilevitz J, Balaban NQ. 2017. TDtest: easy detection of bacterial tolerance and persistence in clinical isolates by a modified disk-diffusion assay. Sci Rep 7:41284.

31. Lazarovits G, Gefen O, Cahanian N, Adler K, Fluss R, Levin-Reisman I, Ronin I, Motro Y, Moran-Gilad J, Balaban NQ, Strahilevitz J. 2022. Prevalence of Antibiotic Tolerance and Risk for Reinfection Among Escherichia coli Bloodstream Isolates: A Prospective Cohort Study. Clin Infect Dis Off Publ Infect Dis Soc Am 75:1706–1713.

32. Lewis K. 2010. Persister cells. Annu Rev Microbiol 64:357–372.

33. Gaca AO, Colomer-Winter C, Lemos JA. 2015. Many means to a common end: the intricacies of (p)ppGpp metabolism and its control of bacterial homeostasis. J Bacteriol 197:1146– 1156.

34. Maslowska KH, Makiela-Dzbenska K, Fijalkowska IJ. 2019. The SOS system: A complex and tightly regulated response to DNA damage. Environ Mol Mutagen 60:368–384.

35. Walker BJ, Abeel T, Shea T, Priest M, Abouelliel A, Sakthikumar S, Cuomo CA, Zeng Q, Wortman J, Young SK, Earl AM. 2014. Pilon: an integrated tool for comprehensive microbial variant detection and genome assembly improvement. PloS One 9:e112963.

36. Aono R, Yamasaki M, Tamura G. 1979. High and selective resistance to mecillinam in adenylate cyclase-deficient or cyclic adenosine 3’,5’-monophosphate receptor protein-deficient mutants of Escherichia coli. J Bacteriol 137:839–845.

37. Zhu K, Zhang Y-M, Rock CO. 2009. Transcriptional regulation of membrane lipid homeostasis in Escherichia coli. J Biol Chem 284:34880–34888.

38. Acosta MBR, Ferreira RCC, Padilla G, Ferreira LCS, Costa SOP. 2000. Altered expression of oligopeptide-binding protein (OppA) and aminoglycoside resistance in laboratory and clinical Escherichia coli strains. J Med Microbiol 49:409–413.

39. Yakhnina AA, Bernhardt TG. 2020. The Tol-Pal system is required for peptidoglycan-cleaving enzymes to complete bacterial cell division. Proc Natl Acad Sci 117:6777–6783.

40. Bichara M, Pelet S, Lambert IB. 2021. Recombinational repair in the absence of holliday junction resolvases in E. coli. Mutat Res 822:111740.

41. Górecka KM, Komorowska W, Nowotny M. 2013. Crystal structure of RuvC resolvase in complex with Holliday junction substrate. Nucleic Acids Res 41:9945–9955.

42. Moya B, Barcelo IM, Cabot G, Torrens G, Palwe S, Joshi P, Umarkar K, Takalkar S, Periasamy H, Bhagwat S, Patel M, Bou G, Oliver A. 2019. In Vitro and In Vivo Activities of β-Lactams in Combination with the Novel β-Lactam Enhancers Zidebactam and WCK 5153 against Multidrug-Resistant Metallo-β-Lactamase-Producing Klebsiella pneumoniae. Antimicrob Agents Chemother 63:e00128–19.

43. Moya B, Barcelo IM, Bhagwat S, Patel M, Bou G, Papp-Wallace KM, Bonomo RA, Oliver A. 2017. WCK 5107 (Zidebactam) and WCK 5153 Are Novel Inhibitors of PBP2 Showing Potent “β-Lactam Enhancer” Activity against Pseudomonas aeruginosa, Including Multidrug-Resistant Metallo-β-Lactamase-Producing High-Risk Clones. Antimicrob Agents Chemother 61:e02529–16.

44. Barriere SL, Gambertoglio JG, Lin ET, Conte JE. 1982. Multiple-dose pharmacokinetics of amdinocillin in healthy volunteers. Antimicrob Agents Chemother 21:54–57.

45. Rodvold KA, Gotfried MH, Chugh R, Gupta M, Patel A, Chavan R, Yeole R, Friedland HD, Bhatia A. 2018. Plasma and Intrapulmonary Concentrations of Cefepime and Zidebactam following Intravenous Administration of WCK 5222 to Healthy Adult Subjects. Antimicrob Agents Chemother 62:e00682–18.

46. Neu HC. 1983. Penicillin-binding proteins and role of amdinocillin in causing bacterial cell death. Am J Med 75:9–20.

47. Joseleau-Petit D, Thévenet D, D’Arl R. 1994. ppGpp concentration, growth without PBP2 activity, and growth-rate control in *Escherichia coli*. Mol Microbiol 13:911–917.

48. Antón DN. 1995. Resistance to mecillinam produced by the co-operative action of mutations affecting iipopolysaccharide, *spoT*, and *cya* or *crp* genes of *Salmonella typhimurium*. Mol Microbiol 16:587–595.

49. Vinella D, D’Ari R, Jaffé A, Bouloc P. 1992. Penicillin binding protein 2 is dispensable in Escherichia coli when ppGpp synthesis is induced. EMBO J 11:1493–1501.

50. Anton BP, Raleigh EA. 2016. Complete Genome Sequence of NEB 5-alpha, a Derivative of Escherichia coli K-12 DH5α. Genome Announc 4:e01245–16.

51. Thulin E, Andersson DI. 2019. Upregulation of PBP1B and LpoB in cysB Mutants Confers Mecillinam (Amdinocillin) Resistance in Escherichia coli. Antimicrob Agents Chemother 63:e00612–19.

52. Ulloa ER, Dillon N, Tsunemoto H, Pogliano J, Sakoulas G, Nizet V. 2019. Avibactam Sensitizes Carbapenem-Resistant NDM-1-Producing Klebsiella pneumoniae to Innate Immune Clearance. J Infect Dis 220:484–493.

53. Cerqueira GC, Earl AM, Ernst CM, Grad YH, Dekker JP, Feldgarden M, Chapman SB, Reis-Cunha JL, Shea TP, Young S, Zeng Q, Delaney ML, Kim D, Peterson EM, O’Brien TF, Ferraro MJ, Hooper DC, Huang SS, Kirby JE, Onderdonk AB, Birren BW, Hung DT, Cosimi LA, Wortman JR, Murphy CI, Hanage WP. 2017. Multi-institute analysis of carbapenem resistance reveals remarkable diversity, unexplained mechanisms, and limited clonal outbreaks. Proc Natl Acad Sci U S A 114:1135–1140.

54. Salamzade R, Manson AL, Walker BJ, Brennan-Krohn T, Worby CJ, Ma P, He LL, Shea TP, Qu J, Chapman SB, Howe W, Young SK, Wurster JI, Delaney ML, Kanjilal S, Onderdonk AB, Bittencourt CE, Gussin GM, Kim D, Peterson EM, Ferraro MJ, Hooper DC, Shenoy ES, Cuomo CA, Cosimi LA, Huang SS, Kirby JE, Pierce VM, Bhattacharyya RP, Earl AM. 2022. Inter-species geographic signatures for tracing horizontal gene transfer and long-term persistence of carbapenem resistance. Genome Med 14:37.

55. CLSI. 2023. Performance Standards for Antimicrobial Susceptibility Testing. 33rd ed. CLSI supplement M100. Clinical and Laboratory Standards Institute.

56. Clinical and Laboratory Standards Institute. 2015. Methods for Dilution Antimicrobial Susceptibility Tests for Bacteria That Grow Aerobically - Tenth Edition: Approved Standard M07-A10. CLSI, Wayne, PA, USA.

57. Brennan-Krohn T, Truelson KA, Smith KP, Kirby JE. 2017. Screening for synergistic activity of antimicrobial combinations against carbapenem-resistant Enterobacteriaceae using inkjet printer-based technology. J Antimicrob Chemother 72:2775–2781.

58. Smith KP, Kirby JE. 2016. Verification of an Automated, Digital Dispensing Platform for At-Will Broth Microdilution-Based Antimicrobial Susceptibility Testing. J Clin Microbiol 54:2288–2293.

59. Chen CY, Nace GW, Irwin PL. 2003. A 6 x 6 drop plate method for simultaneous colony counting and MPN enumeration of Campylobacter jejuni, Listeria monocytogenes, and Escherichia coli. J Microbiol Methods 55:475–479.

60. Hall BG, Acar H, Nandipati A, Barlow M. 2014. Growth rates made easy. Mol Biol Evol 31:232–238.

61. Bolger AM, Lohse M, Usadel B. 2014. Trimmomatic: a flexible trimmer for Illumina sequence data. Bioinforma Oxf Engl 30:2114–2120.

62. Li H, Durbin R. 2009. Fast and accurate short read alignment with Burrows-Wheeler transform. Bioinforma Oxf Engl 25:1754–1760.

63. Zuluaga AF, Salazar BE, Rodriguez CA, Zapata AX, Agudelo M, Vesga O. 2006. Neutropenia induced in outbred mice by a simplified low-dose cyclophosphamide regimen: characterization and applicability to diverse experimental models of infectious diseases. BMC Infect Dis 6:55.

64. Kang AD, Smith KP, Berg AH, Truelson KA, Eliopoulos GM, McCoy C, Kirby JE. 2018. Efficacy of Apramycin against Multidrug-Resistant Acinetobacter baumannii in the Murine Neutropenic Thigh Model. Antimicrob Agents Chemother 62:e02585–17.

65. Andes D, Craig WA. 2002. Animal model pharmacokinetics and pharmacodynamics: a critical review. Int J Antimicrob Agents 19:261–268.

66. Endimiani A, Hujer KM, Hujer AM, Pulse ME, Weiss WJ, Bonomo RA. 2011. Evaluation of ceftazidime and NXL104 in two murine models of infection due to KPC-producing Klebsiella pneumoniae. Antimicrob Agents Chemother 55:82–85.

